# Development of a Next Generation SNP Genotyping Array for Wheat

**DOI:** 10.1101/2023.11.27.568448

**Authors:** Amanda J. Burridge, Mark Winfield, Alexandra Przewieslik-Allen, Keith J. Edwards, Imteaz Siddique, Ruth Barral-Arca, Simon Griffiths, Shifeng Cheng, Zejian Huang, Cong Feng, Susanne Dreisigacker, Alison R. Bentley, Gina Brown-Guedira, Gary L. Barker

## Abstract

High throughput genotyping arrays have provided a cost effective, reliable and interoperable system for genotyping hexaploid wheat and its related germplasm pool. Existing, highly cited arrays including our 35K Axiom Wheat Breeder’s genotyping array and the Illumina 90K iSelect array were designed based on a limited amount of varietal sequence diversity and with imperfect knowledge of SNP positions. Recent progress in sequencing wheat varieties and landraces has given us access to a vast pool of SNP diversity, whilst technological improvements in array design has allowed us to fit significantly more probes onto a 384-well format Axiom array than was previously possible. Here we describe a novel High Density Axiom genotyping array, the *Triticum aestivum* Next Generation array (TaNG), largely derived from whole genome skim sequencing of 204 elite wheat lines and 111 wheat landraces taken from the Watkins “Core Collection”. We use a novel “minimal marker” optimisation approach to select up to six SNPs in each 1.5 MB region of the wheat genome with the highest combined varietal discrimination potential. A design iteration step allowed us to test and replace skim-sequence derived SNPs which failed to convert to reliable Axiom markers, resulting in a final design, designated TaNG1.1 with 43,372 SNPs derived from a haplotype-optimised combination of novel SNPs, DArTAG-derived and legacy wheat Axiom markers. We show that this design has an even distribution of SNPs across chromosomes and sub-genomes compared to previous arrays and can be used to generate genetic maps with a significantly higher number of distinct bins than our previous Axiom array. We also demonstrate the improved performance of TaNG1.1 for Genome Wide Association Studies (GWAS) and its utility for Copy Number Variation (CNV) analysis. The array is commercially available, and the marker annotations, initial genotyping results and software used to generate the optimised marker sets are freely available.

## Introduction

Single Nucleotide Polymorphism genotyping arrays (SNP arrays) play an important role in advancing studies of genetic variations in both animal (Chen *et al*., 2014a) and plant populations (Bassil *et al*.,2015; Koning-Boucoiran *et al*.,2015; van Geest *et al*., 2017). They allow the identification, and analysis of up to hundreds of thousands of SNPs in a single assay (see review by You *et al*., 2018) and provide a powerful platform for highlighting genome-wide, sequence variability between individuals in and between populations. Thus, SNP genotyping arrays provide a high-throughput and cost-effective way to analyse genetic diversity and have been widely used to generate genetic linkage maps, study evolutionary relationships, unravel functional genomics and support conservation efforts. They have also proved to be highly valuable in breeding programmes being used for marker assisted selection (MAS) (Thompson, 2014; Arruda *et al*., 2016), genome-wide association studies (GWAS) (McCouch et al., 2016; Negro *et al*., 2019; Balagué-Dobón *et al*., 2022; Yu *et al*., 2023), and the mapping of QTL (Xu *et al*., 2017; Stadlmeier *et al*., 2018). Their power and utility are evidenced by the large number of arrays available for crop species (strawberry - Verma *et al*., 2017; rose - Koning-Boucoiran *et al*., 2015; chrysanthemum - van Geest *et al*., 2017; potato-Vos *et al*., 2015; rice - Chen *et al*., 2014b; Kim *et al*., 2022; Deware *et al*., 2023; maize - Unterseer *et al*., 2014).

Bread Wheat is a crop species of vital importance to the world economy providing. Genotyping arrays have played, and continue to play, a critical role in the genotyping of hexaploid bread wheat. The use of genotyping arrays has allowed researchers to rapidly genotype wheat varieties, identify genetic variants associated with important traits, and develop markers for use in breeding programmes. In particular, the development of high-density genotyping arrays has enabled researchers to genotype large numbers of wheat samples and identify genetic variants with a high degree of accuracy. For wheat, several SNP arrays have been developed (Wang *et* al., 2014; Winfield *et al*., 2016; Allen *et al*., 2017; Rimbert *et al*., 2018; Soleimani *et al*., 2020; Sun *et al*., 2020). These arrays contain a large number of SNPs and have been demonstrated to be effective tools for linkage analysis, QTL mapping of important traits and genome-wide association analysis (Allen *et al*., 2017; Bourke *et al*., 2018; Vukosavljev *et al*.,2016).

However, previously developed genotyping arrays for wheat suffer from uneven marker distribution and marker redundancy due to linkage disequilibrium (LD). Uneven marker distribution means that regions of the genome being over- or underrepresented. This can lead to bias in the results and limit the ability to accurately detect genetic variants in certain regions of the genome. This is particularly problematic for bread wheat, which has a large, hexaploid complex genome with significant structural variation. In addition to these technical limitations, older genotyping arrays are also limited by the genetic diversity of the populations used to develop them. Bread wheat is a highly diverse crop with significant genetic variation both between and within different populations.

Therefore, genotyping arrays developed using a limited set of wheat lines may not capture the full range of genetic diversity present in the crop. As a consequence of these limitations, scientists and breeders have called for a new generation of wheat genotyping arrays that overcome these technical and biological challenges, provide a more comprehensive view of the genome, and capture the full range of genetic diversity present in bread wheat’s pangenome. Here, we describe the development of a new SNP genotyping array for wheat, the TaNG Array, that has been designed to overcome several of these issues and, thus, provide a more comprehensive coverage of the genome than previous versions.

## Methods

### Marker selection: Skim Sequence Sourced Probes

The SNP calls generated from skim sequence data from 315 wheat accessions (204 elite wheat lines and 111 wheat landraces taken from the Watkins “Core Collection” - Supplementary File S1) were used as the source of SNPs for haplotype optimisation (Cheng *et al*. 2023). Varieties with >= 1% heterozygous loci were excluded. SNPs were initially filtered to have a maximum of 0.5% heterozygous calls among all varieties, a minimum minor allele frequency of 0.01, a minimum call rate of 0.95 and a minimum mapping quality score of 5000. SNPs with a flanking sequence mapping to more than one genome location in the IWGSC v1.0 Chinese Spring genome assembly using the BWA version 0.7.12-r1039 were removed. Additionally, SNPs were checked by BLAST (blastn v2.6.0+) against the IWGSC v1.0 Chinese Spring genome assembly and those matching multiple locations were excluded. Each chromosome was then divided into 1.5 Mb intervals and up to six SNPs representing the highest combined discriminatory power were selected for each interval (Winfield *et al*., 2020). The haplotype optimisation pipeline is available at https://github.com/pr0kary0te/GenomeWideSNP-development.

### Existing Marker Designs

For cross compatibility, SNPs for which there are existing markers from various platforms were also included in the design. That is, 2,528 markers selected for trait association or physical locations were taken from the CIMMYT Wheat 3.9K DArTAG array (https://excellenceinbreeding.org/toolbox/services/mid-density-genotyping-service) as were 4,220 of the best performing markers from the existing Axiom™ Wheat Breeder’s Genotyping Array (Allen *et al*., 2017). In addition, a public call was made to researchers and wheat breeders to nominate markers from existing arrays which they would like to see included on the TaNG array; this call resulted in 1,223 marker nominations (Supplementary File S3, *v1.1 Probe Details*). The final design also has 936 cross-platform probes with the now discontinued Illumina 90k iSelect array (Wang *et al.,* 2014) and 8,232 cross-platform probes with the Wheat 660k Axiom array (Sun *et al.,* 2020).

On an initial screening of an early version of the array (designated TaNG v1.0) against a diverse set of 119 elite and 60 landrace accessions, 16,507 SNPs failed to convert to polymorphic SNP assays. These markers were replaced with 14,774 selected from the Axiom™ Wheat HD Genotyping Array (Winfield *et al*. 2016), to maximise the differentiation of varieties described above. This final, optimised array design, designated TaNG v1.1 (Thermo Fisher catalogue number 551498), contains 43,373 markers (Table 1; Supplementary File S3, *TaNG v1.1 Marker Details*.) Some markers may be present on multiple arrays under different names, when this is the case, pseudonyms are given in each column.

**Table 1:**
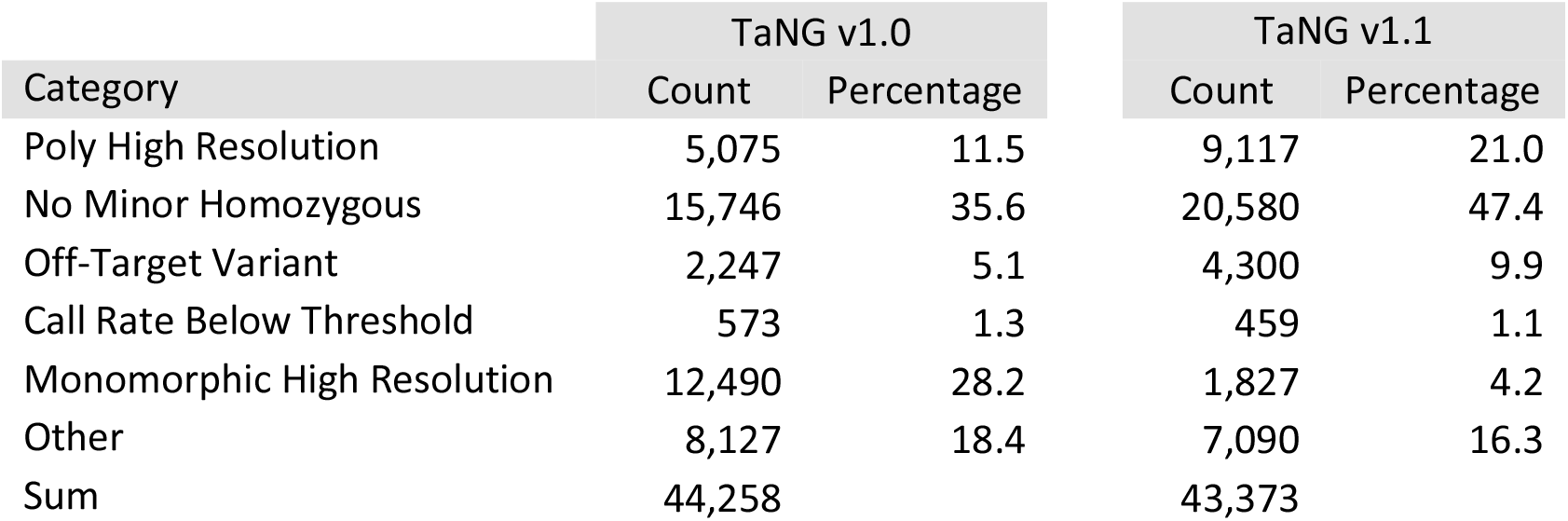
Marker quality categories for TaNG v1.0 and TaNG v1.1.

### Modification of priors

For consistency, the same Dish-QC probes were used as the 35k Axiom™ wheat breeder’s genotyping array for generation of the non-genotype producing DQC sample quality metrics. For probe quality, of the markers on TaNG v1.1, 299 generated clusters that were not correctly identified during allele calling using Applied Biosystems’ software package Axiom™ Analysis Suite v5.2.0.65. To ensure the correct genotype call for these alleles, the analysis file was modified with sequence specific priors for the affected markers (’SSP’ in Supplementary S3, *Probe Details*). The modified analysis file designated ‘Axiom_TaNG1_1.r4’ is available from the Thermo Fisher website.

### Genotyping

Genomic DNA from wheat leaf tissue 14 day after germination was prepared as described in Burridge *et al*. (2017) for samples listed in Supplementary File S4. Genotyping was performed using 11ul of 25ng ul-1 DNA in water. Array processing was performed using the GeneTitan system according to the procedure outlined in Axiom™ 2.0 Assay 384HT Array Format Automated Workflow User Guide (Applied Biosystems). Allele calling was performed using Applied Biosystems’ software package Axiom™ Analysis Suite v5.2.0.65 using prior file Axiom_TaNG_SNP.r1 for the first array design (v1.0) and Axiom_TaNG1_1.r4 for the final array design (v1.1). In all cases a Dish QC of 0.8 for *T. aestivum (Initial Testing, USA Material)* datasets and 0.6 for the wild relative genotyping. A sample QC call rate of 80% and 75% was used for *T. aestivum* and wild relative sets respectively. The SNP QC cut-off for ‘Call Rate Below Threshold’ was 95%. Comparisons of technical replication was made using markers across all probe quality categories with ‘No-call’ genotypes omitted from comparison. The TaNG v1.1 Array is available from Thermo Fisher Scientific with catalogue number 551498.

### Copy Number Variation (CNV)

The CNV analysis and Manhattan plots were generated for all accessions screened on TaNG v1.1 using Axiom™ Analysis Suite v5.2.0.65, with prior file Axiom_TaNG1_1.r4 and the annotation file Axiom_TaNG1_1.r4.annot.db. No samples were excluded from reference creation. The recommended minimum base lengths and probe numbers for each CNV state were followed from Axiom™ Copy Number Data Analysis Guide (r3 May 2022, MAN0026736).

### SNP Effect Predictions

SNP effect predictions were made using the Variant Effect Predictor (VEP) hosted on the EnsemblPlants website (http://plants.ensembl.org/Triticum_aestivum/Tools/VEP). The genome selected was that of *Triticum aestivum*. Variant call format (vcf) files were uploaded to the website and the web tool run using default settings.

### Genetic Map Construction

For the three mapping populations, markers with more than 10% missing data were removed. The remaining markers were tested for significant segregation distortion using a chi-square test. The software program MapDisto v. 1.7 (Lorieux, 2012) was used to assemble the loci into linkage groups using likelihood odds (LOD) ratios with a LOD threshold of 6.0 and a maximum recombination frequency threshold of 0.4. Linkage groups were ordered using the likelihoods of different locus-order possibilities and the iterative error removal function (maximum threshold for error probability 0.05) in MapDisto. The Kosambi mapping function (Kosambi, 1944) was used to calculate map distances (cM) from recombination frequency. Maps were drawn in MapDisto with bins represented by a single marker.

### Consensus Chromosome Assignment

Where possible, markers were assigned to a chromosome based on consensus of calls from four different data sets: i) physical position from BLASTing sequence to IWGSC assembly v1.0; ii) Avalon x Cadenza genetic map (10,113 markers); iii) Apogee x Paragon map (4,673 markers); iv) Oakley x Gatsby map (7,733 markers). To be assigned a consensus chromosome, the calls from at least two of the data sets had to agree; if a marker had only a physical position or only conflicting calls, it was not assigned a consensus. A comparison was made between the consensus calls and initial physical call to estimate agreement (Supplementary File S3); this analysis was performed taking into account the origin of the markers, skim sequence derived vs acquired from earlier genotyping platforms (820K Array, 35K Array, DArT).

### Genome Wide Association Study

The GWAS analysis was performed using the Watkins collection accessions and associated phenotype data as described in (Cheng *et al.,* 2023) to compare the core 10M SNPs from the sequenced dataset (Cheng *et al.,* 2023); SNPs from the Axiom™ Wheat Breeder’s Genotyping Array (CerealsDB) and those of the TaNG Array v1.1. Extreme outlier values of phenotypic data were removed. Kinship matrix was calculated as the covariate using GEMMA-kin. Based on these, GWAS was performed using GEMMA (v0.98.1) with parameters (gemma-0.98.1-linux-static -miss 0.9 - gk kinship.txt) and gemma-0.98.1-linux-static -miss 0.9 −lmm -k kinship.txt). In-house R scripts were used to visualize the results.

## Results

### Marker testing

The SNP markers selected for inclusion on the array were from skim sequence data from 315 wheat accessions (204 elites and 111 landraces – Supplementary File S1) and from existing genotyping arrays (see Methods). Markers taken from the Axiom™ Wheat HD Genotyping Array (Winfield *et al*. 2016) - hereafter referred to as the 820K Axiom™ Wheat HD Genotyping Array - and the Axiom™ Wheat Breeder’s Genotyping Array (Allen *et al*., 2017) - hereafter referred to as the 35K Array - were entirely exonic, while those derived from sequence were intronic, exonic and intergenic. As a panel, the markers were evenly distributed throughout the genome based on positions relative to IWGSC RefSeq v1.0 (International Wheat Genome Sequencing Consortium *et al.,* 2018).

The initial array design (v1.0) was screened using a standard collection of 182 elite cultivars and landraces (Supplementary File S4, *Sample Details and Genotyping*, sheet ’TaNG v1.0 Genotyping’). The sample call rate ranged from 94.5% to 98.8%. Based on their cluster patterns, probes were classified into the following six categories: Poly High Resolution; No Minor Homozygous; Off-Target Variant; Call Rate Below Threshold; Monomorphic High Resolution and Other (Figure 1; Table 1). The first three categories are considered most useful as they generate accurate polymorphic genotype calls. Of the 44,258 probes on the initial array, 23,068 (52%) fell into the three useful categories (Table 1). Approximately 28% were monomorphic; that is, they generated a strong signal to indicate the target sequence was present, but no polymorphism was detected. Of these monomorphic markers, 93% were derived from skim-sequence data rather than sourced from existing genotyping arrays, indicating that these SNPs failed to convert into useful Axiom markers.

**Figure 1.**
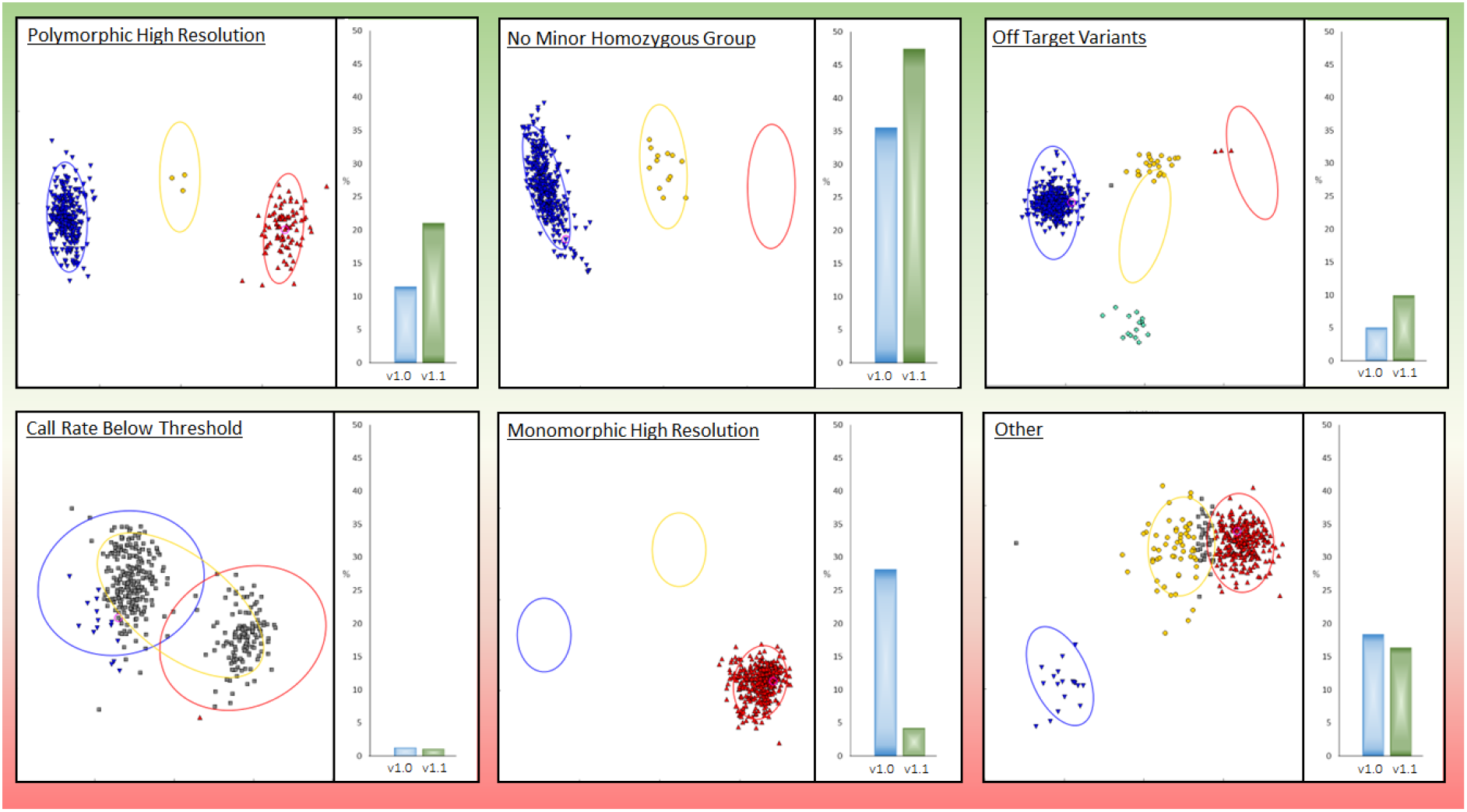
The percentage of probes in each probe quality category by array type. TaNG v1.1 has an increased ratio of the ‘high quality’ categories (Polymorphic High Resolution, No Minor Homozygous Group and Off-Target Variants) and a decreased ratio of probes in the ‘low quality’ categories (Call Rate below Threshold, Monomorphic High Resolution and Other).

Of the accessions used for screening TaNG Array v1.0, 144 were also present in the original skim sequencing panel (Supplementary File S1) thus allowing direct comparison to be made between genotype calls on the two platforms. Given the sequencing-derived genotypes for the 144 varieties, only four of the 12,490 markers reporting monomorphisms were predicted to be monomorphic, and only 112 of these apparently monomorphic markers were expected to have less than 10 instances of the minor allele.

### Design optimisation

Due to the large number of monomorphic probes on the first iteration of the TaNG array (version 1.0), the array was redesigned. Monomorphic probes were replaced with probes from the Axiom™ Wheat HD Genotyping Array proven to be polymorphic; these replacement probes were selected by re-running the marker optimisation algorithm with monomorphic markers excluded from the input file, whilst other markers were retained as they had performed well in screening (Supplementary File S3 – *Probe Details*). Additional markers were integrated into the optimised design by analysing combining genotyping data from our existing 820K Wheat HD Genotyping Array with that derived from TaNG v1.0, where the same samples had been run on both platforms. This improved array, designated TaNG v1.1, was screened against an extended collection of elite cultivars, landraces and other *Triticum* accessions (Supplementary File S4, *Sample Details and Genotyping,* sheet ’TaNG v1.1 Genotyping’). The sample call rate ranged from 84% to 99.8%. Compared to the initial implementation of the array, TaNG v1.1 showed an increased number of markers in each of the useful probe quality categories and a decreased number in each of the less useful categories (Table 1). Therefore, all further study was based on TaNG v1.1 and, thus, from this point forward, all results and discussion refer to comparisons between the 35K Array version v1.1 of the new array.

### Marker Distribution Across Chromosomes (Physical Designation)

The TaNG v1.1 Array has more markers in total than the 35K Array (43,373 vs 35,143) and, for all chromosomes except 1D and 2D, there are more markers assigned to each chromosome (Table 2). Furthermore, markers are more evenly distributed between the 21 wheat chromosomes and the number of markers per chromosome better reflects chromosome size (Supplementary Figure S5 - *Marker Distribution,* sheet *’Markers per Chromosome’*).

**Table 2.**
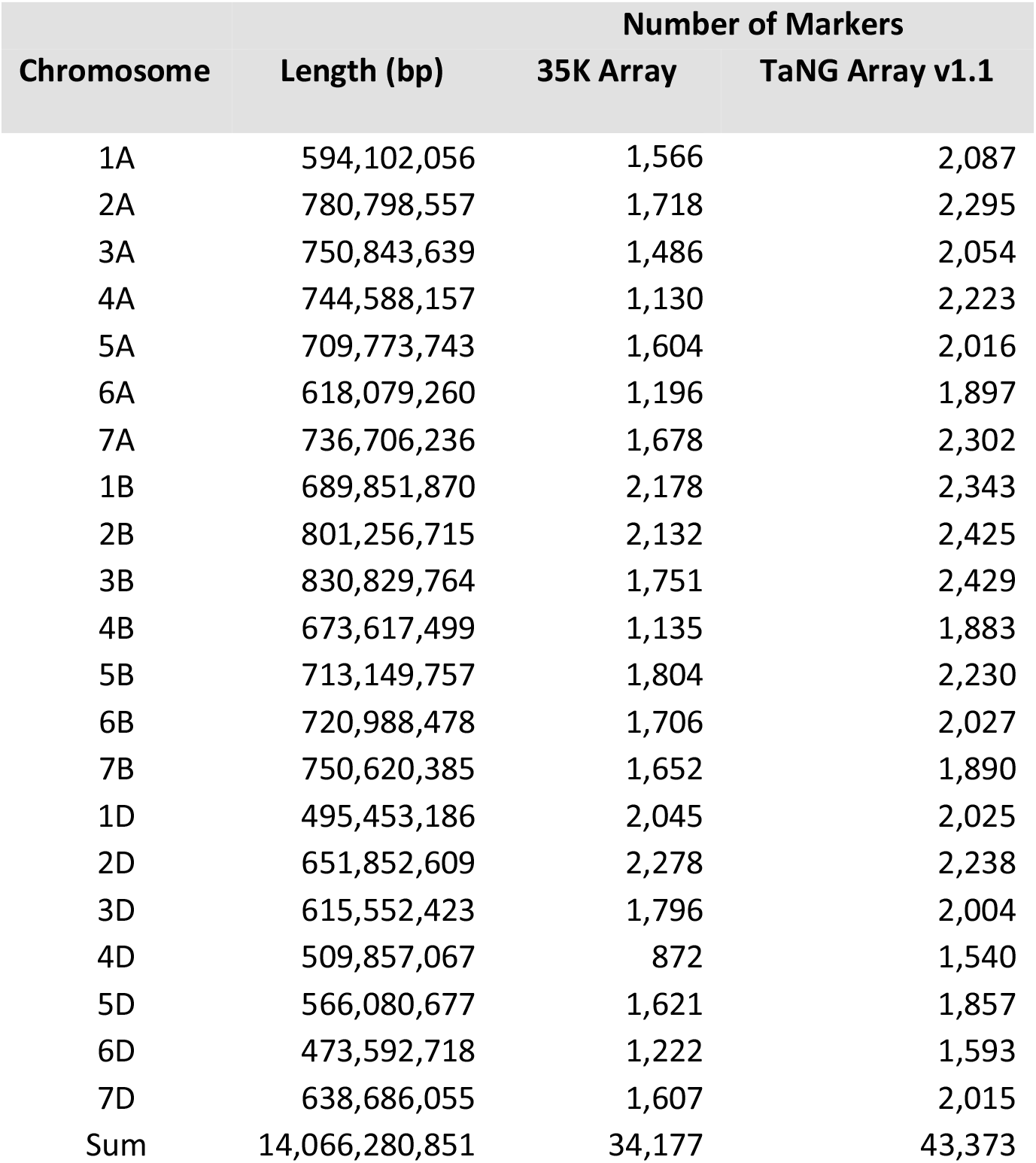
Chromosome lengths in variety Chinese Spring (based on IWGSC v1.0 assembly) versus number of markers on the 35K and TaNG v 1.1 Arrays.

| Chromosome | Length (bp) | Number of Markers |  |
| --- | --- | --- | --- |
|  |  | 35K Array | TaNG Array v1.1 |
| 1A | 594,102,056 | 1,566 | 2,087 |
| 2A | 780,798,557 | 1,718 | 2,295 |
| 3A | 750,843,639 | 1,486 | 2,054 |
| 4A | 744,588,157 | 1,130 | 2,223 |
| 5A | 709,773,743 | 1,604 | 2,016 |
| 6A | 618,079,260 | 1,196 | 1,897 |
| 7A | 736,706,236 | 1,678 | 2,302 |
| 1B | 689,851,870 | 2,178 | 2,343 |
| 2B | 801,256,715 | 2,132 | 2,425 |
| 3B | 830,829,764 | 1,751 | 2,429 |
| 4B | 673,617,499 | 1,135 | 1,883 |
| 5B | 713,149,757 | 1,804 | 2,230 |
| 6B | 720,988,478 | 1,706 | 2,027 |
| 7B | 750,620,385 | 1,652 | 1,890 |
| 1D | 495,453,186 | 2,045 | 2,025 |
| 2D | 651,852,609 | 2,278 | 2,238 |
| 3D | 615,552,423 | 1,796 | 2,004 |
| 4D | 509,857,067 | 872 | 1,540 |
| 5D | 566,080,677 | 1,621 | 1,857 |
| 6D | 473,592,718 | 1,222 | 1,593 |
| 7D | 638,686,055 | 1,607 | 2,015 |
| Sum | 14,066,280,851 | 34,177 | 43,373 |

For broad scale distribution of markers, the chromosomes, regardless of their reported lengths (Table 2) were divided into 20 equally sized bins and the number of markers in each bin totalled and plotted (Figure 2A); markers on TaNG v1.1 are more evenly distributed across the chromosomes than those on the 35K Array. That is, there is neither a bias in marker number towards the telomeres nor a relative paucity of markers across the centromeres. At a smaller scale, marker distribution was determined by dividing chromosomes into 10 Mb bins and counting the number of markers in each (Figure 2B). The number of markers in the 10 Mb bins is much less variable and there are no extreme outliers with very low or very high numbers of markers. For example, on the 35K Array, the region 240 - 290 Mb on chromosome 4A is represented by only 6 markers; on TaNG v1.1 the same region is represented by 125 markers (Supplementary File S5 - *TaNG1-1*). Indeed, on chromosomes 3A and 4A the 35K Array has no markers assigned at all to a small number of 10 Mb bins (Supplementary File S5 - *Marker Distribution*, sheet ’*Distribution (20 Equal Bins)’*).

**Figure 2.**
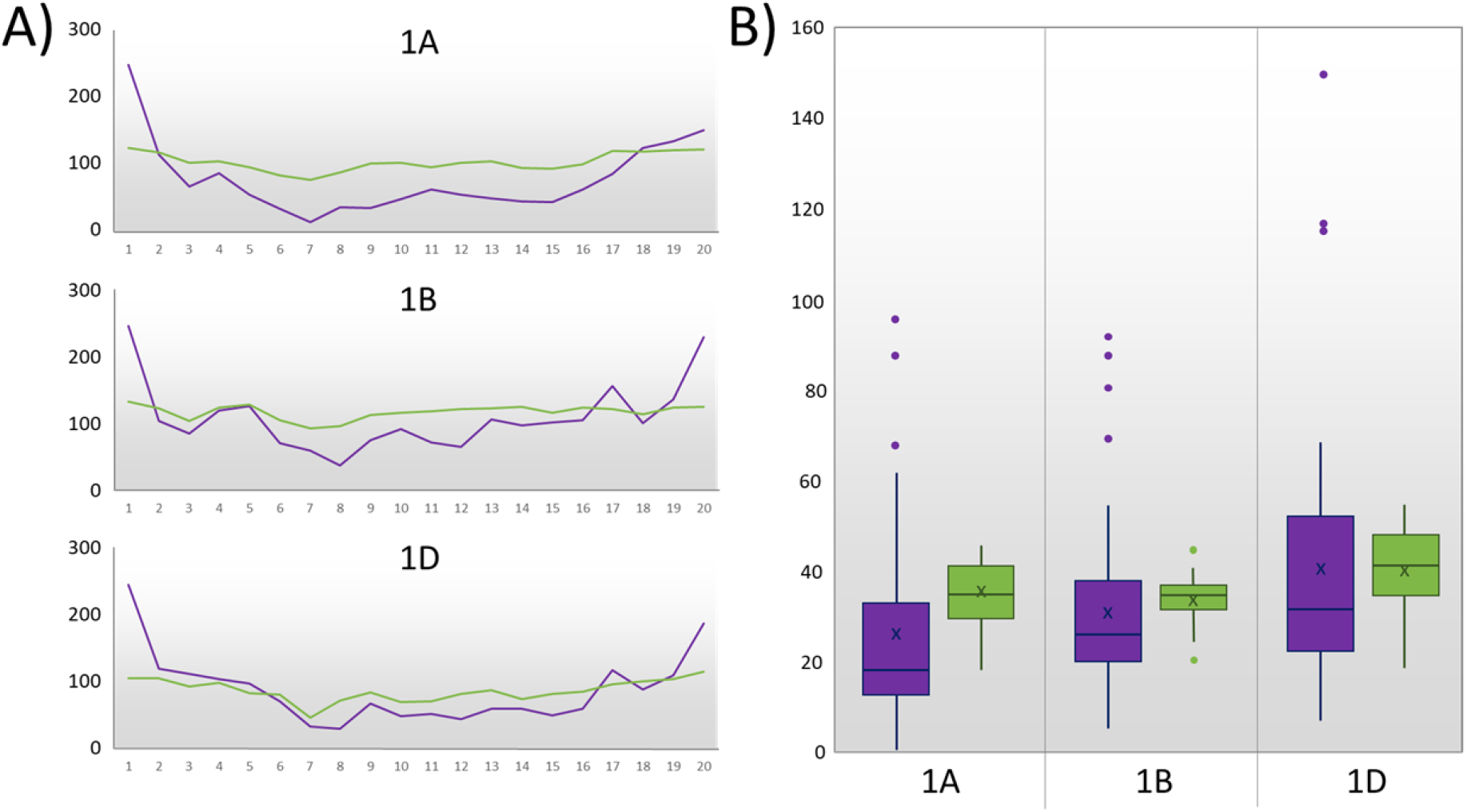
The physical distribution of chromosome 1 markers on the TaNG v1.1 (green) and 35K Wheat Breeder’s Arrays (purple). A) Number of markers in each of 20 bins spanning the chromosome. B) Box and whisker plots of the number of markers per 10 Mb bin across the chromosome. There is a greater number of markers on the TaNG v1.1 Array (green boxes) and these are more evenly distributed than on the 35K Array (purple boxes). See Supplementary File S5 for plots of all chromosomes.

Although there is a relatively strong relationship between chromosome length and the number of markers assigned to that chromosome, the density of markers is relatively higher on the smaller chromosomes than on the larger chromosomes: mean number of markers per 1 Mb on the A, B and D genome is 3.0, 2.9 and 3.4, respectively.

### Source of SNP variation

The older 35K Array SNPs were derived from exome capture sequencing, based on a set of genes de novo assembled genes from cDNA sequencing of Chines Spring (Winfield *et al*. 2012).

Unsurprisingly, a variant effect prediction analysis annotated over 86% of these SNPs to be within or immediately adjacent to coding regions (Figure 3). In contrast, more than 27% of the SNPs on the new TaNG1.1 array were annotated as having an intergenic origin (Figure 3).

**Figure 3.**
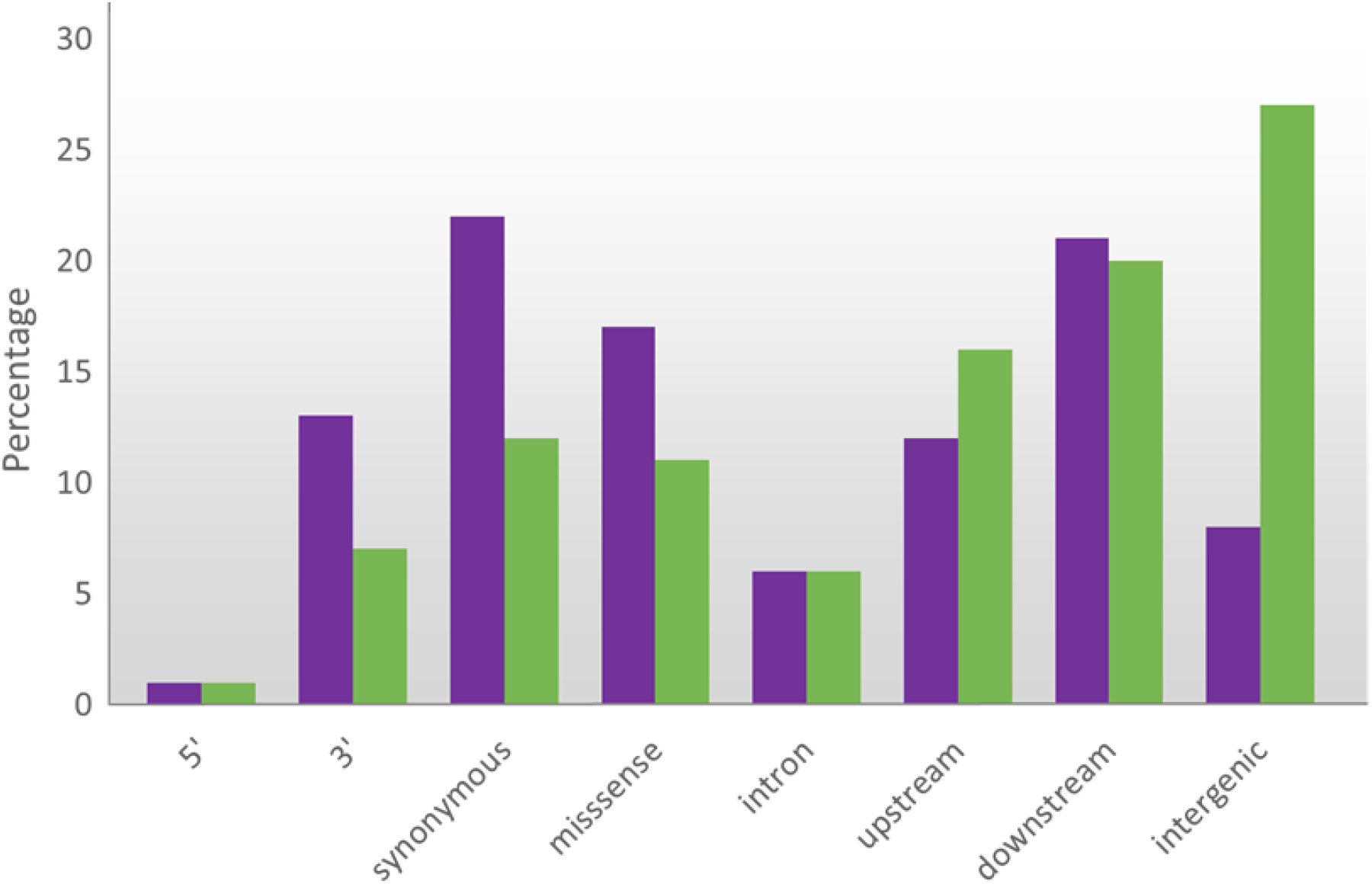
SNP variant effect for the 35K Axiom™ Wheat Breeder’s genotyping array (purple boxes) and the TaNG v1.1 Array (green boxes) derived from the Variant Effect Predictor (VEP) web tool at EnsemblPlants (Release 110; Fergal et al., 2023) with Triticum aestivum selected as genome. Given that the markers on the 35K Array were derived from exome capture libraries, there are few SNPs from intergenic regions.

### Replicate Testing for Reproducibility

As a measure of reproducibility between technical replicates, the accessions ‘Paragon’ and ‘Cadenza’ were genotyped as four aliquots from the same DNA extraction (Supplementary File S4 - *Sample Details and Genotyping*). The genotype correlation was extremely high with 99.86% and 99.49% correlation for ‘Paragon’ and ‘Cadenza’ respectively. This represents a less that 1% technical error rate. The genotyping errors were predominantly a mis-call between the homozygous (AA, BB) and heterozygous (AB) states with only two and four probes presenting a change in homozygous call (‘hom-hom mis-call’) in ‘Paragon’ and ‘Cadenza’ respectively. Only six markers presented a genotyping error between both accessions, each on a different chromosome suggesting that the 1% error rate was random in nature (Supplementary File S4 - *Sample Details and Genotyping*).

### Genetic Map Construction

The genetic location of markers, and performance of the haplotype optimisation marker selection method was tested by generating genetic maps using three mapping populations. These were the Avalon x Cadenza (AxC) and Oakley x Gatsby (OxG) double haploid populations, and Apogee x Paragon (AxP) produced by single seed descent to the F5 generation. For the three maps, AxC, OxG and AxP, 10,113, 7,734 and 4,673 markers were assigned to linkage groups, respectively (Supplementary File S6).

The AxC and AxP populations had previously been genotyped using the Axiom™ Wheat Breeder’s Genotyping Array, making it possible to compare the position of markers on the arrays. The TaNG AxC genetic map consisted of 10,113 markers with 1,652 unique locations, while the Wheat Breeder’s AxC genetic map consisted of 7,237 markers with 1,082 unique locations (Figure 4, Supplementary File 6). The TaNG AxP genetic map consisted of 4,673 markers with 1,984 unique locations, while the Wheat Breeder’s AxC genetic map consisted of only 2,997 markers with 1,519 unique locations. The TaNG array both increased the number of markers and the number of unique positions for both the AxC and AxP genetic maps. For all chromosomes the number of SNPs and the chromosome length (cM) was greater using the TaNG v1.1 array than the maps previously constructed using data from the Axiom 35k Wheat Breeder’s Genotyping Array. Although restricted by the limits of recombination of the population, the new array gives a greater number of more evenly spaced markers for all three populations.

**Figure 4.**
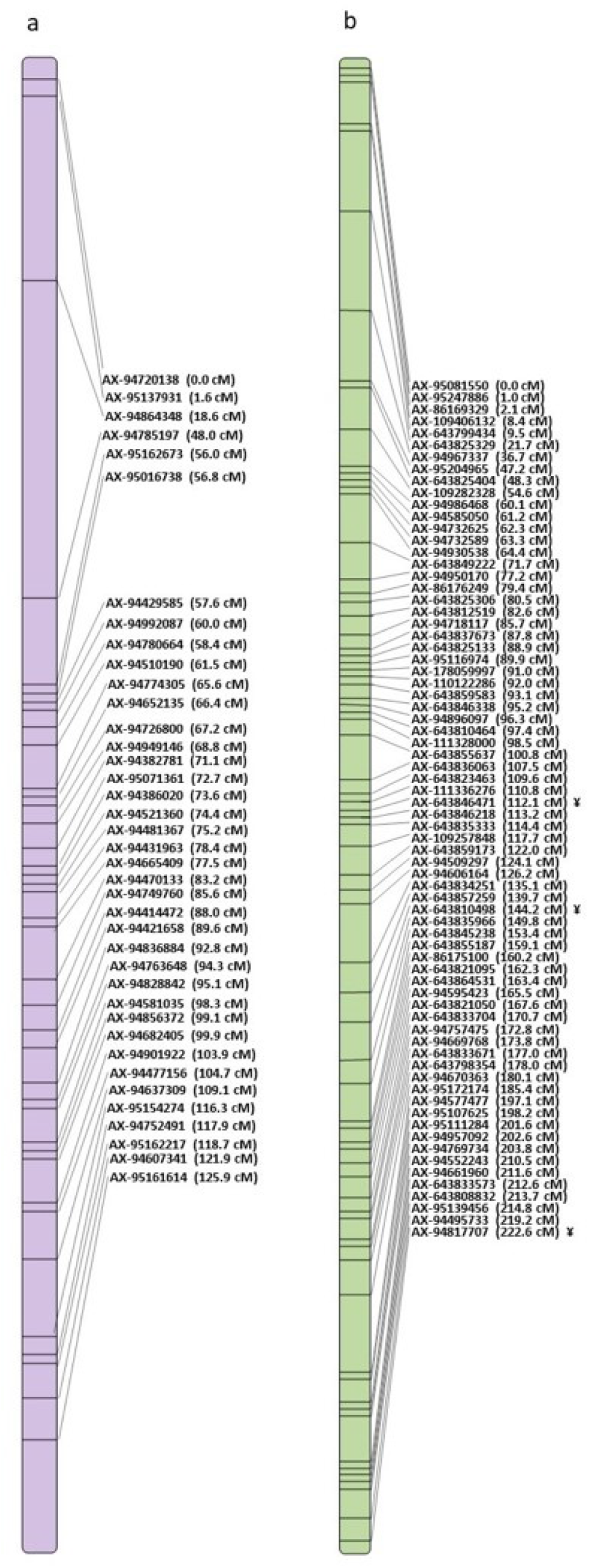
Comparison of Avalon x Cadenza genetic maps for chromosome 1A using (a) 35K Array map data and (b) TaNG v1.1 map data, showing the distribution and density of markers.

The genetic map positions of markers from all three genetic maps were compared to the physical assignment based upon alignment of sequences to the Chinese Spring, IWGSC v1.0 reference assembly: physical assignment based on alignment to IWGSC assembly v1.0; AxC map; AxP map; OxG map. A marker was assigned a consensus chromosome only when at least two of the assignments were the same. In total, 12,981 markers were assigned a consensus chromosome (Supplementary File S3).

The comparison showed that the physical assignment based on skim sequence data was close to 100% accurate whilst that derived from previous platforms was less reliable, especially for D-genome markers (Figure 5). There was good correlation between physical position of the markers and the cM position of the bin to which they were assigned. However, physical assignment for markers taken from previous platforms (820K Array, 35K Array and DArT marker) were more variable with an average concordance of 85% for A and B genome markers but as low as only 40% for D genome markers.

**Figure 5.**
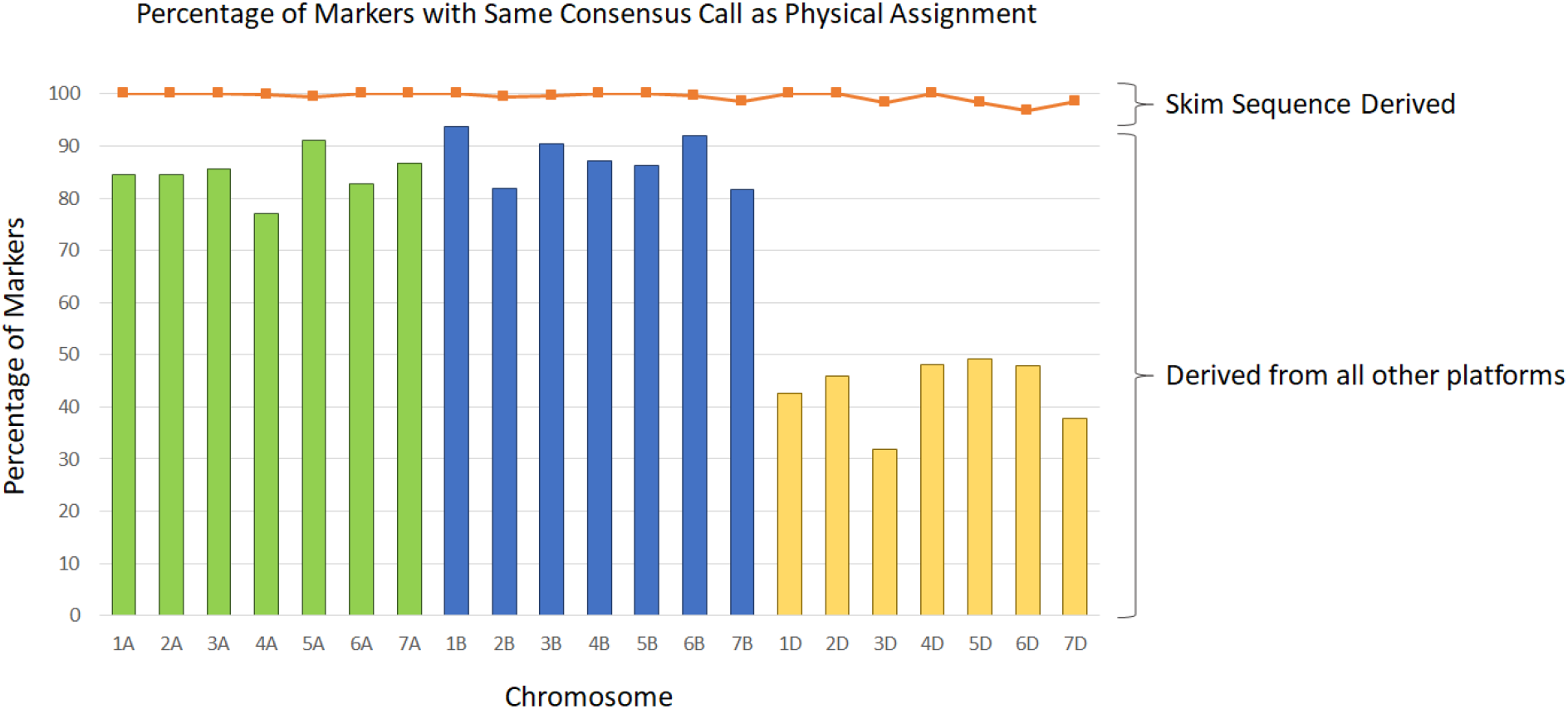
Comparison of chromosome location for markers based on physical and consensus assignment. Only markers with a consensus chromosome assignment are shown. Markers with physical positions derived from previous genotyping platforms (820K Array, 35K Array and DArT markers) are represented by bars. Markers derived from skim sequence data are represented by a line plot.

### CNV Analysis

Copy number events were observed across each chromosome with the exception of 4D, 5A and 5D (Figure 6; Supplementary S7 – *CNV summary Table*). The number of CNV varied per accession, some reported none while 12 were detected in the *T. aestivum* accession ‘Captor’ and 23 in the wild relative *‘T. kiharae’.* The length of detected CNV regions ranged from 8.7 Mb to 780 Mb with more reduced CNV losses than gains. Multiple regions are present with common variations across the screened accessions. Large regions of variance are present on 2A (Loss: 479,492 – 24,414,560 in 50 accessions; Gain: 202,646,955 – 358,094,640 in 36 accessions), 5B (Gain: 149,551,898 – 249,788,187 in 32 accessions; Loss: 490,000,000 - 520,000,000 in 50 accessions) and 1B (Gain and Loss: 1,203,929 – 123,969,113) (Figure 6; Supplementary file S7).

**Figure 6:**
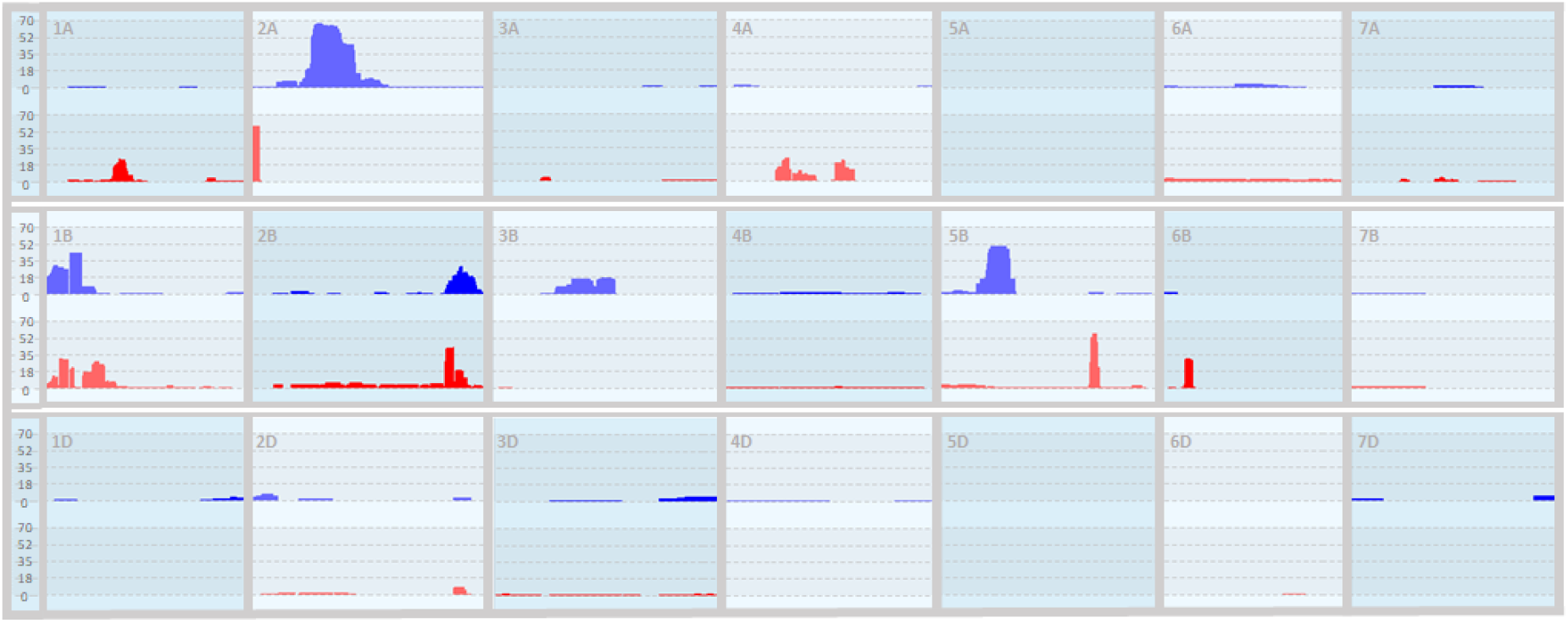
Copy Number Variance (CNV) frequency histogram for all samples genotyped with the TaNG v1.1 array in the initial screening (Supplementary S4) across all chromosomes. Regions of copy number gain are displayed in the top track (blue) and regions of copy number loss are displayed on the bottom track (red) for each chromosome.

### Diverse Material Screening

As the initial array screening was performed with a mostly European collection of elite cultivars and landraces (Supplementary File S4), additional genotyping was also performed using different sources of material. A collection of USA material grouped by geographic origin was genotyped as the initial screening panel (Supplementary File S4). The marker call rate was consistently high with 42,476 markers (98%) generating a call in 90% of samples. The performance category of markers was also high with only 3,746 probes (8.6%) found to be monomorphic across the dataset. The genotype data clearly distinguished the USA material by region (Figure 7) with a clear separation of North region and East region germplasm.

**Figure 7:**
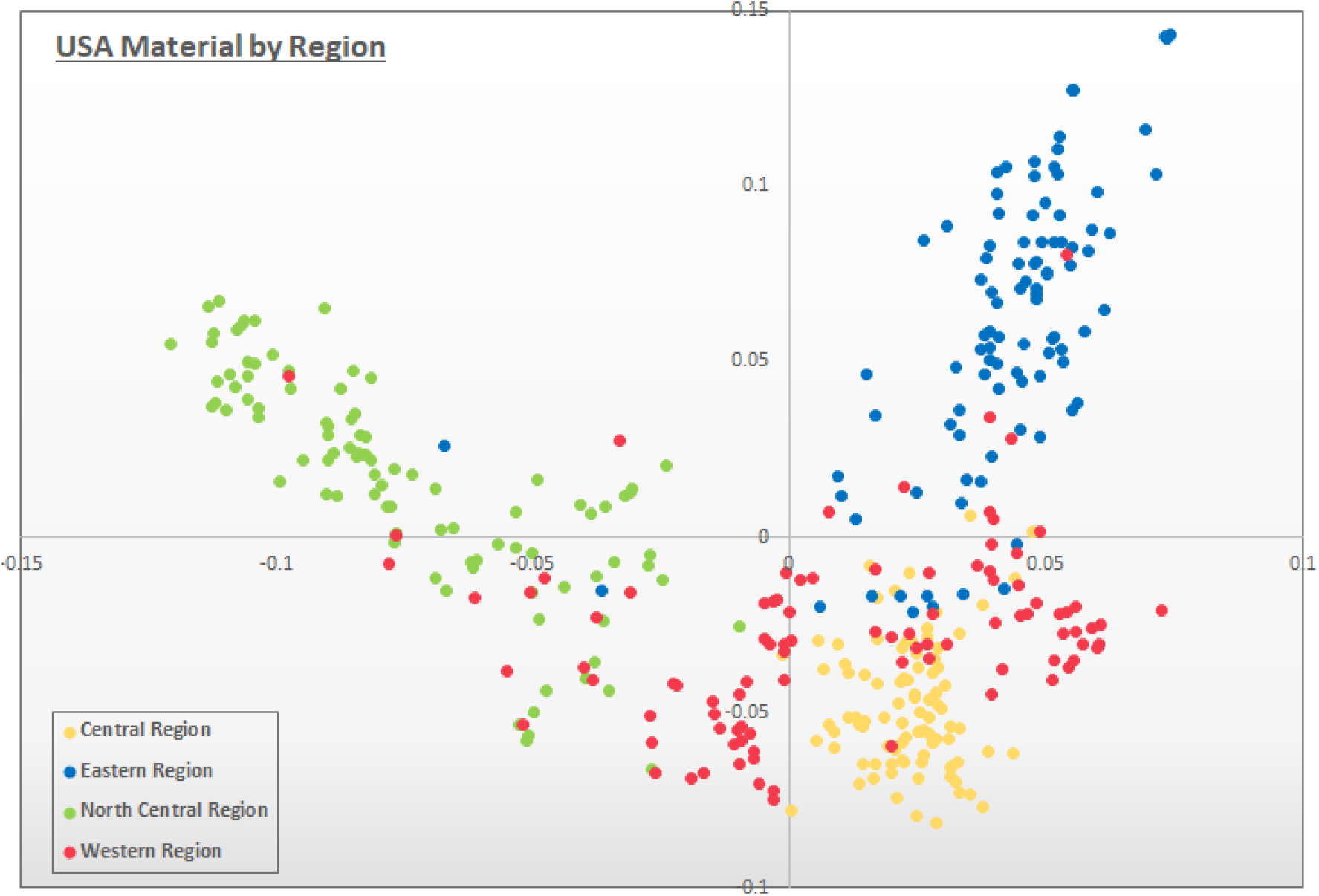
Principle Component Analysis plot based on the USA material collections coloured by donating region. Variation in PC1 and PC1 is 5.05 and 3.93 respectively. Source data is available in Supplementary File S4.

A collection of 81 wheat wild relatives including *Aegilops, Amblyopyrum, Secale, Thinopyrum* and other *Triticum* species were also genotyped alongside 50 Durum (*T. Turgidum ssp. durum*) landrace accessions (Supplementary File S4) to examine the suitability of the array for genotyping pre-breeding wild relative material alongside *T. aestivum*. While not all samples could hybridise, 34,588 markers (80%) generated a genotype call across at least 90% of the samples. The majority of markers clustered unclearly with the ‘Other’ performance category (18,784; 43%) but very few markers were monomorphic (4,008; 9.2%).

### GWAS Analysis

As a test of performance for the optimised SNP selection, three traits were selected for GWAS analysis: Heading Date; Response to Leaf Rust; Response to Stem Rust. While the previous 35k Breeders array was unable to identify a significant QTL for any of these traits, the 43K TaNG v1.1 array was able to identify QTL that favourably compared to the entire 10 million SNP panel generated from whole genome sequencing (Figure 8).

**Figure 8:**
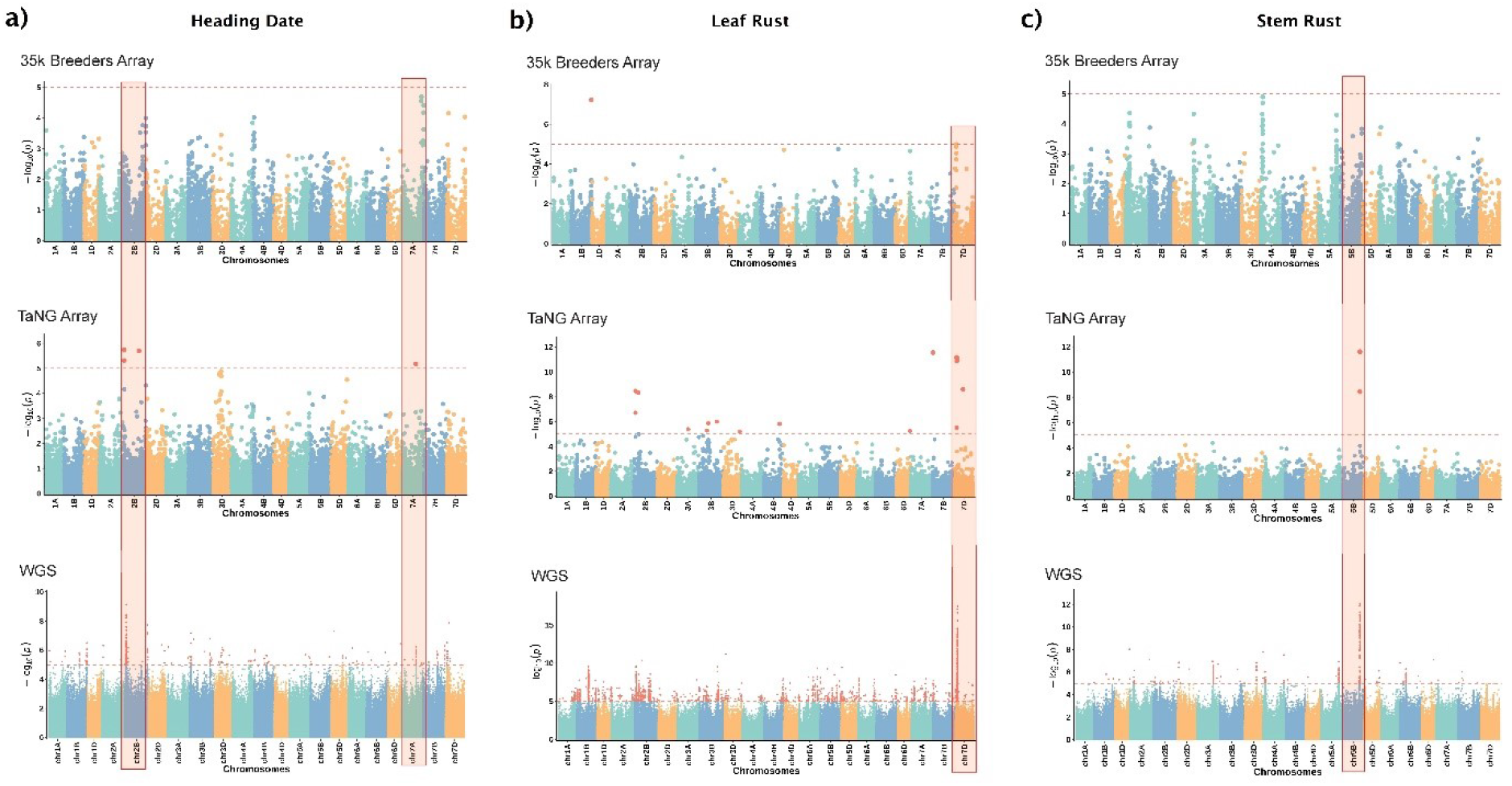
Genome Wides Association Study (GWAS) using the previous 35k Breeders Array, the new TaNG v1.1 array and the 10 million SNPs detected in sequenced data for three traits. A) Heading date B) Leaf Rust C) Stem Rust. Identified QTL are highlighted in red.

## Discussion

### Array Justification

The existing 35K Axiom™ wheat breeder’s genotyping array designed in 2011 has been widely used by academics and breeding companies with 288 citations in research areas recently spanning pathogen resistance (Nannuru *et al*. 2022, Grover *et al*. 2022), yield (Sheoran *et al*. 2022), grain nutrient quality (Rathan *et al*. 2022) and grain architecture (Kumari *et al*. 2023). While a valuable tool, improvements to technology and our understanding has made it possible to improve upon the design in several ways. The most noticeable difference is the source of SNPs. In the 820k Axiom™ Wheat HD Genotyping Array (Winfield et al., 2016), the 35K Axiom™ wheat breeder’s genotyping array (Allen *et al*., 2017) and the 90k wheat iSelect array (Wang *et al*., 2014) used exome-capture sequences as the source of putative SNPs. This resulted in all SNPs being within or very close to genes, resulting in uneven chromosomal distribution. Uneven marker distribution means that some regions of the genome are overrepresented, while others are underrepresented. Furthermore, many arrays focused only on European breeding lines as a source of material narrowing the pool of gene-based SNPs even further. Such focussed selection of samples coupled with the tendency to used sequence capture prior to sequencing could lead to the exclusion of rare alleles and a strong ascertainment bias (You *et al*., 2018). Recent large scale skim sequencing (Cheng *et al*. 2023) of a highly diverse set of globally significant breeding and landrace accessions made available a new source of SNPs free of ascertainment bias and suitable for global users.

Prior the release of the IWGSC Chinese Spring reference (IWGSC, 2018), genetic maps were used to obtain marker location information. In the 35k Array this led to significant ascertainment bias due to prioritizing SNPs that could be placed on the Avalon x Cadenza or Rialto x Savanna maps (Allen *et al*., 2017). For SNPs without polymorphisms between these parents, the locations (physical or genetic) were initially unknown. After the release of the CS reference, the distribution was found to be variable, chromosomes had markers clustered together in some regions with large gaps in between with no marker coverage. In the areas of marker clusters, many would be in linkage disequilibrium (LD) meaning that the SNPs would be inherited together more often than expected by chance. LD represent a duplication of effort, as additional markers are no more informative. With a gold standard genome sequence now available and the physical positions of all SNPs now known, more careful consideration can be made with regards to the marker distribution.

### Two-Step Design Process for Improved Performance

The original draft TaNG v1.0 array design showed promising performance for many of the markers but with 12,490 (28.2%) markers that were found to be monomorphic. While the ‘Call Rate Below Threshold’ and ‘Other’ categories may be considered less useful categories, the designation of these categories depends on the samples used. Probes with an ‘Other’ category in one sample set may have a clear genotype clustering in another sample set. However, as probes were designed to be polymorphic within the test set, monomorphic calls were an indication of marker design failure. This high-level of marker failure is not unexpected when converting sequence-derived SNPs to markers in a polyploid and was observed with our first high wheat axiom array (Winfield *et al*. 2016) Because consistently monomorphic or failed probes are of no value, a two-step approach of screening followed by re-design was used. The final design (TaNG v1.1) was produced by combining genotyping results from our original 820K Axiom™ Wheat HD Genotyping Array (Winfield *et al*., 2016) with those from TaNG v1.0. The SNP optimisation algorithm used in the initial design was re-applied to this combined dataset for marker selection to ensure that the replacement markers were fully integrated with the new design. The resulting TaNG v1.1 array had a decreased ratio of probes in all of the ‘low quality’ categories (Figure 1) and an improvement in D genome coverage (Supplementary File S5).

The use of a two-step design method has been employed in the design of other genotyping arrays such as maize (Unterseer *et al*., 2014) and pear (Montanari *et al*., 2019) to produce a high quality and reproducible array design. The number of monomorphic probes on the initial testing of the TaNG v1.1 array was 4.2%, lower than other commercial Axiom arrays such as Pine (10.1%; Perry *et al*., 2020), Groundnut (23.7% monomorphic; Pandey et al., 2017), Chickpea Array (45.7%; Roorkiwal *et al*., 2018) and the original 35K Array (4.5%; Allen *et al*., 2017). As not all of the accessions used for marker generation were used in the array testing (Supplementary Files S1 and S4), this value may be lower when additional diverse lines are used.

### Haplotype Optimisation to Combine New and Existing Probe Designs

The bias towards SNPs in genic regions in the previous 820K HD Array and 35K Array resulted in markers being negatively correlated with chromosome length; that is, relative to length there were more markers on the shorter chromosomes than on the longer ones. On the TaNG v 1.1 Array there is a strong positive correlation which in addition to physical distribution, has been optimised for haplotype grouping. We employed a novel selection algorithm to select the optimal combination of SNPs in each 1.5 Mb bin of each wheat chromosome. Rather than allocating the same number of SNPs to each bin, the SNPs within a bin are minimally correlated with each other to avoid effective duplication. In this way, each SNP has considerably more diagnostic power than those identified at random. The result was a more even physical distribution than the 35K Array and greater diagnostic power even when fewer markers are present per bin (Figure 2). The power of this method was illustrated in use of the array in GWAS (Figure 8). Previously, genotyping arrays have been limited for GWAS applications due to the limited marker density. Even in regions that appear to have good coverage, SNPs in LD provide redundant information about the same genetic variation and bias kinship estimations.

To compliment the sequence derived SNPs, markers selected from existing arrays by haplotype optimisation or public nomination were included. The incorporation of a subset of markers from existing genotyping arrays can maintain continuity and consistency when comparing genetic data across different studies and populations. Combined datasets can enhance statistical power and increase the ability to detect genetic associations without the regeneration of data. In the case of wheat breeding, multi-year studies are not unusual and benefit from the use of consistent marker sets even as genotyping technologies evolve. This approach is already established in medical genotyping arrays such as the Transplant array (Li *et al*., 2015), Axiom Asia Precision Medicine Research Array and the Axiom Human Genotyping SARs-COV-2 Array (Thermo Fisher Scientific) which all contain cross-platform markers. More recently, agricultural genotyping arrays such as the Axiom 50K 4Tree array, Axiom 44K Rice and Infinium Apple arrays (Guilbaud et al., 2020; Affymetrix Datasheet P/N GGNO05960 Rev. 1; Howard et al., 2021) have been designed to include markers compatible with previous edition genotyping arrays.

As the previous 35k Array used exome-capture derived sequences for SNP discovery, there were far fewer intergenic SNPs included than has been possible with the TaNG v1.1 array (Figure 3). For SNPs contained within genes it has been possible to use existing information to identify 157 which are associated with important traits (Supplementary File S3).

### Array Features

The TaNG v1.1 array has a stable 1% technical variation which is in line with other Axiom arrays (GenomeWide 6.0 Human array; Hong *et al*., 2012) and genotyping technologies such as the 1% variation reported using SNP DArTSeq (Nantongo *et al*., 2022; Alam *et al*., 2018), 0.5% reported using SeqSNP (Harper *et al*., 2020) and the 0-1% variation reported using Infinium (Senthilvel *et al*., 2019; Cai *et al*., 2017; Pavy *et al*., 2016). The nature of the genotyping errors between technical replicates were predominantly hom-het mis-calls. This may be due to the probe partially binding to a secondary homoeologous site, sample contamination or due to difficulties in the genotype calling software in identifying a clear cluster. As all assays have been test-screened with a diverse set of accessions (Supplementary File S4) any probes which presented difficulties in genotype calling have been assigned SNP specific priors as described in the methods to ensure consistent calling by the software (Supplementary File S3) making the calling of SNPs on the array as accurate as possible.

Genetic maps were constructed using the TaNG v1.1 array and compared to the 35K Array. On average markers were more evenly distributed across the chromosome with a higher number of unique locations represented and less markers clustered together at the same location (Figure 4, Supplementary File S6). Although limited by recombination in the populations this represents a significant improvement in the resolution of the maps and utility of the markers for accurately mapping QTLs and marker assisted selection. In addition, a comparison of genetic map position and physical position on the Chinese Spring v1.0 reference sequence allowed an analysis of the accuracy of the physical position assignment. This revealed the physical assignment based on skim sequence data is close to 100% accurate whilst that derived from previous platforms is less reliable, especially for D-genome markers (Figure 5). To aid correct placement of markers, the mapping locations are included in Supplementary File S3.

The ability of a genotyping array to perform Copy Number evaluations is limited compared to sequencing methods, but the ease of use and the high-throughput nature allows for insight into sample panels and populations. We used copy number variation (CNV) analysis to characterise the accessions screened. Several common regions of increased (CNV gain) or reduced (CNV loss) signal were observed which could potentially represent deletions, introgressions or repeat regions. Some of these regions are already well documented such as the 1RS introgression from rye on 1BS which is reported to result in variable copy number (Xiong *et al*., 2023) and the *Ae. ventricosa introgression* on 2A (Gao *et al.,* 2021) which is commonly found in wheat due to the addition of the Lr27 resistance gene. Deletions were also clearly represented by a copy number loss in the genotyping data such as the ph1 deletion on 5B (Figure 6: 5B).

### Supporting Data

The bread wheat genome already contains significant genetic variation and much work is being done to enhance the germplasm with novel alleles from wide crosses. The previous 35k Breeders array had previously been used with wheat wild relative material in a pre-breeding context (Kumar *et al.,* 2020; Wright *et al.,* 2023) and for elite durum wheat cultivars (Kabbaj *et al.,* 2017; Ganugi *et al.,* 2021; Shewry *et al.,* 2023). The genotype calls generated on the TaNG v1.1 array across a diverse set of wheat relative material here (Supplementary File S4) illustrate that secondary and tertiary genepool material may also be genotyped alongside *T. aestivum* accessions. As the primary purpose of the array was the genotyping of *T. aestivum*, we suggest a DQC cut off of 0.6 to be used for wheat relative material to account for the absence of some reference sequences used to generate the DQC metric. The grouping of diploid, tetraploid and hexaploid material with a wide range of ancestral genomes created a valuable insight the relatives for which the TaNG array may successfully hybridise but for more accurate genotyping study, we suggest that samples be grouped by project before genotype calling.

To support cross-platform projects and better support data sharing, the TaNG array has incorporated SNP probes from other public arrays such as the CIMMYT Wheat 3.9K DArTAG array and the previous 820k HD Wheat array and 35k Breeders array. Further to this, other commercial genotyping platforms have included our probes in the same way. We have compiled these including the 90k iSelect array (Wang *et al.,* 2014) and 660k array (Cui *et al.,* 2017) synonyms with full sequences and known trait associations from literature (Supplementary File S3). We believe that together with the TaNG array, this will be valuable resource for all researchers working across genotyping platforms.

In essence, SNP genotyping arrays have revolutionised the way researchers and breeders study plant genetics and manipulate traits. They provide a high-throughput and cost-effective way to analyse the genetic makeup of plant populations, enabling more targeted and efficient research and breeding efforts. As technology continues to advance, SNP genotyping arrays will continue to be a cornerstone of plant science, contributing to sustainable agriculture, crop security, and our understanding of plant biology.

## Acknowledgements

The array processing was performed by the Bristol Genomics Facility.

We are grateful to the Wheat Genetic Improvement Network for making public the mapping data relating to the Avalon x Cadenza population. This population of doubled-haploid (DH) individuals was developed by Clare Ellerbrook, Liz Sayers and the late Tony Worland (John Innes Centre), as part of a Defra funded project led by ADAS. The parents, having contrasting canopy architectures, were originally chosen by Steve Parker (CSL), Tony Worland and Darren Lovell (Rothamsted Research).

We also express gratitude to the Germplasm Resources Unit (GRU) at the John Innes Centre (JIC) for much of the elite, landrace and wheat relative germplasm used to screen the array, to the USDA Germplasm Resource Information Network (GRIN) for much of the USA germplasm and to the NBRP-Wheat gene bank for additional wild relative germplasm.

